# Inositolphosphate glycans and a fucosylated xyloglucan oligosaccharide are accumulated upon *Arabidopsis thaliana/ Botrytis cinerea* infection

**DOI:** 10.1101/2021.06.25.449971

**Authors:** Aline Voxeur, Julien Sechet, Samantha Vernhettes

## Abstract

In mammals, insulin is involved in controlling blood glucose levels and its role in modulating immunity is being more and more documented. This hormone promotes the release of inositolphosphate glycans (IPG) which act as mediators. In plants, one IG has already been identified in plant culture cells (Smith and Fry, 1999; Smith et al., 1999) but, to our knowledge, no IPG have been yet identified. Here, we discovered 7 IPG that are accumulated upon *Arabidopsis thaliana-Botrytis cinerea* interaction, concomitantly with oligogalacturonides and a fucosylated xyloglucan oligosaccharide. Further structural characterization showed that they come from the hydrolysis of polar heads of Serie A to H glycosylinositol phosphorylceramides presumably via a phospholipase C activity. Taken together with the emerging role of insulin as immune regulator, these results question the role of IPG as damage associated molecular pattern both in animal and plant kingdoms.

## Introduction

Oligosaccharins are biological active oligosaccharide^1^. Their biological activities include the stimulation of skotomorphogenesis^2^, growth promotion^3-6^ and anti-auxin activity^7-8^. We also know for decades that oligogalacturonides (OGs), oligomers of •-1,4-linked galacturonic acid trigger plant defense response when they are exogenously applied^9-11^. Until very recently^12^, almost none of the oligosaccharins studied have been detected *in planta*, thus questioning their role and their identity *in planta*. Here we identified unknown oligosaccharides accumulated during *Arabidopsis thaliana-Botrytis cinerea interaction*. While we rather expected to characterize cell wall derived oligosaccharides, we discovered that several inositol phosphate glycans (IPG) deriving from plant glycosylinositol phosphorylceramides (GIPCs) are highly accumulated upon infection.

First, using high-performance size-exclusion chromatography (HP-SEC) coupled with high resolution mass spectrometry (HRMS)–based method in negative mode, we have set up a strategy aimed at identifying putative oligosaccharides accumulated over time upon *Botrytis cinerea* infection. We selected the main ions detected upon infection and increasing continuously between 12 and 18 hours post-infection with a mass-to-charge ratio (*m /z*) superior to 263 that corresponds to a dipentose (**Fig. 1A**). This leads to the identification of 36 candidates included 16 OGs, 4 being oxydized (**Fig. 1B**). Amongst the 18 non-OG candidates, we discarded 5 members for which we did not obtain confident elemental compositions. Finally, this workflow resulted in 13 candidates. 9 are phosphorylated and 3 do not present any elemental composition compatible with an oligosaccharide structure (**Fig. 1C**). We observed that 8 of them were also detected in cell wall water extract of non-infected plants.

**Figure 1.**
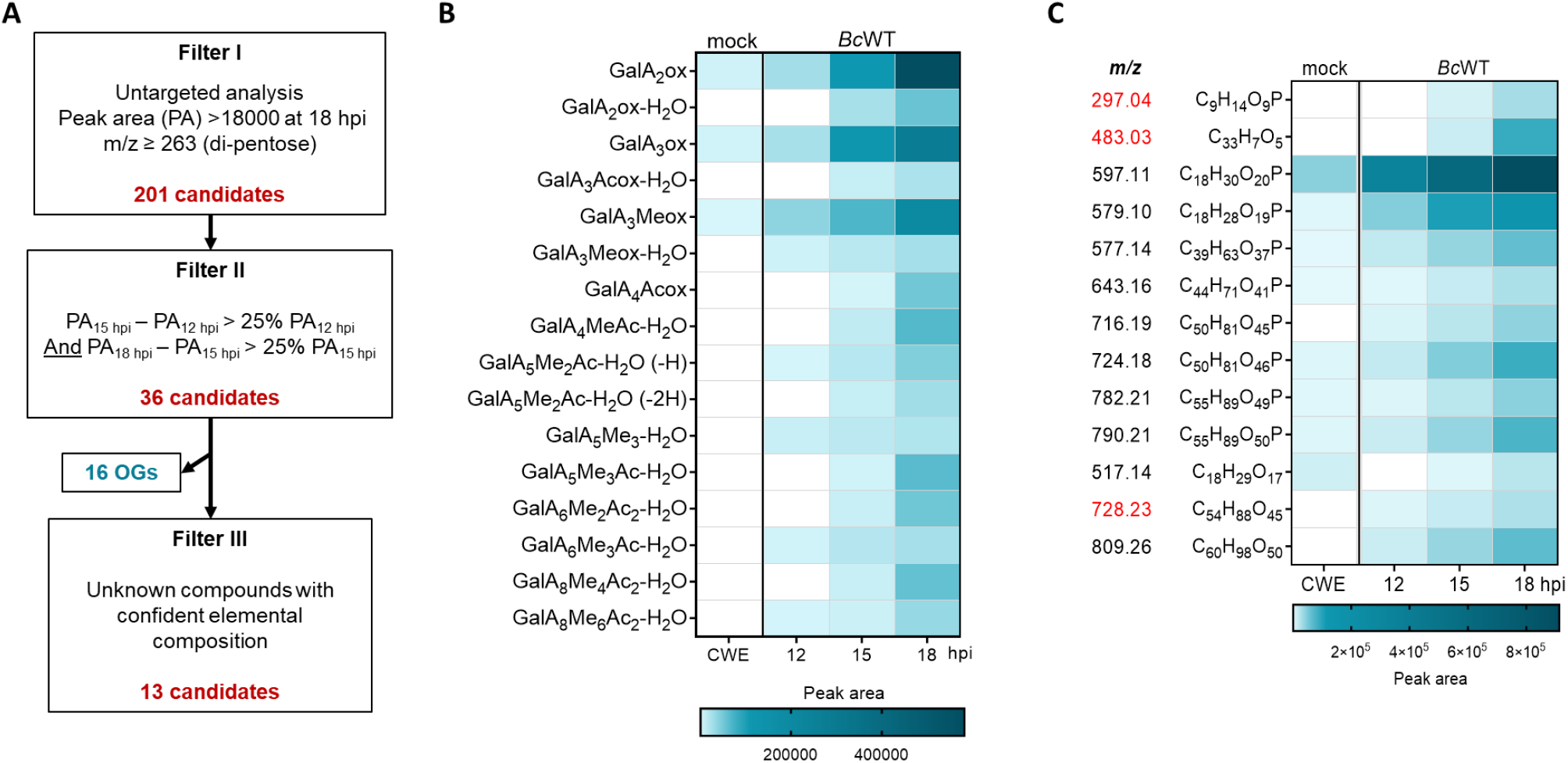
Phosphorylated oligosaccharides are accumulated upon *Arabidopsis thaliana-Botrytis cinerea* infection. **A)** Flow chart of the 3-step filtering strategy used to select putative oligosaccharides accumulated upon infection. **B**) Heat map displaying the oligogalacturonides (OG) accumulated over time upon infection. **C**) Heat map displaying the non OG candidates accumulated over time upon infection. Candidates in red do not have any elemental compositions compatible with an oligosaccharide structure. PA: Peak area; hpi: hours post infection; OG: oligogalacturonides; CWE: Cell wall water extract; BcWT: *B. cinerea* wild type strain; GalA: Galacturonic acid; Me: Methylester group; Ac: Acetylester group; OGs are named GalA_x_Me_y_Ac_z_. Subscript numbers indicate the DP and the number of methyl- and acetylester groups, respectively.

We first focused our study on the most abundant candidate at *m / z* 597 [M - H]^-^ for which the software predicted the formula C_18_ H_30_ O_20_ P. This compound coelutes with the *m / z* 579 ion (predicted formula C_18_ H_29_ O_19_ P^-^) which we attributed to the formation of a [M-H-H_2_ O]^-^ anion. The *m / z* 597 MS^2^ spectrum displays a *m /z* 97 ion corresponding to H_2_ PO_4_ ^-^ that comforts the predicted formula (**Fig. 2A**). The fragment ion at *m /z* 373 is likely formed from the decarboxylation of the *m /z* 417 ion, the latter corresponding to the loss of a hexose (Hex) moiety from the parental ion [M-H-180]^-^. The ion at *m /z* 241 was next assigned the loss of an uronic acid (UA) from the *m / z* 373. Last, the *m / z* 259 and 241 ions disgnostic of phosphorylated inositol (Ino-P)^13^ suggest that *m / z* 597 ion corresponds to an inositolphosphate glycan (IPGs) composed of a phosphorylated inositol, an uronic acid and a hexose (**Fig. 2B**). We next fragmented the candidate at *m / z* 517 (C_18_ H_29_ O _17_^-^) (**Fig. 2C**). We observed fragment ions at *m /z* 161 and 179 corresponding to hexosyl or inositol residue. We also detected characteristic fragment ions of uronic acids at *m / z* 175 [M-H_2_ O-H]^-^ and 113 [M-COO]^-^. Consistent with the *m / z* 597 fragmentation mechanism, the fragment ion at *m / z* 293 is formed by the loss of a hexose or an inositol and a further decarboxylation. We conclude that *m /z* 517 likely corresponds to the non-phosphorylated form of the *m /z* 597 candidate.

**Figure 2.**
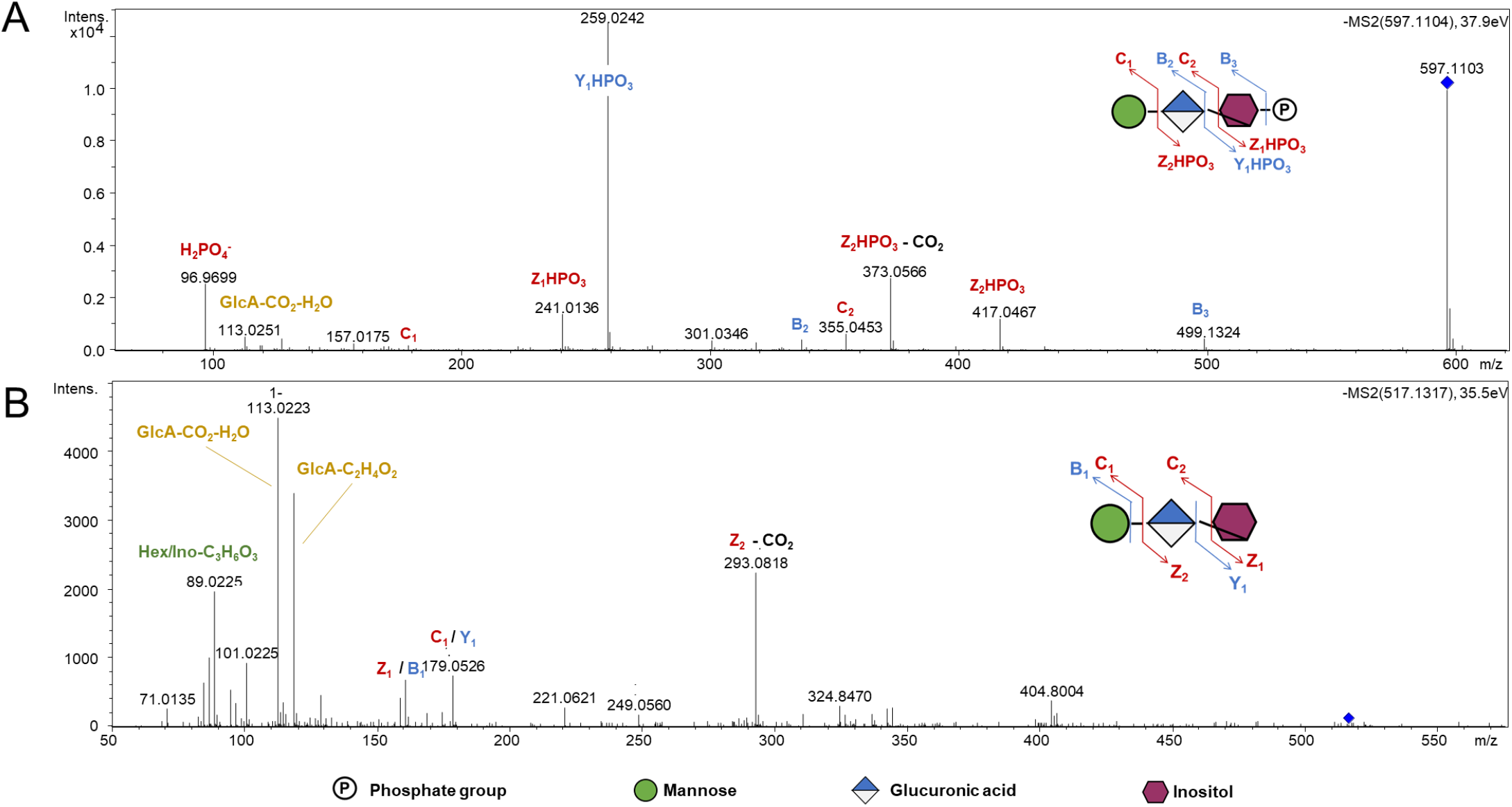
Short Inositol(Phosphate) Glycans (GIPs) are accumulated upon *Arabidopsis thaliana-Botrytis cinerea* infection. **A**) MS^2^ fragmentation pattern of *m/z* 597 in negative mode. **B**) MS^2^ fragmentation pattern of *m/z* 517 in negative mode. GlcA: Glucuronic acid, Intens.: signal intensity.

The *m / z* 259 (Ino/Hex-P) and 373 (UA-Ino/Hex-P) ions observed in *m / z* 597 MS^2^ spectrum were found in the fragmentation pattern of six doubly charged candidates at *m /z* 577 (C_39_ H_63_ O _37_P^2-^), 643 (C_14_ H_71_ O_41_ P^2-^), 716 (C_50_ H_81_ O_45_ P^2-^), 724 (C_50_ H_81_ O_46_ P^2-^), 782 (C_55_ H_89_ O_50_ P^2-^), and 790 (C_55_ H_89_ O_50_ P^2-^). According to the fragmentation pattern obtained from *m / z* 577 and 643 (**Fig. 3A-D**), these molecules contain one additional hexose and three and four additional pentoses respectively, likely linked to a Hex-UA-Ino/Hex-P core. The cross-ring fragments at *m /z* 353, 485, 617 and 749 (B_4•_ + C_2_ H_4_ O_2_) might be produced by the cleavage of an inositol or a hexose residue. We favor the first hypothesis since no fragment corresponding to three or four pentoses plus two hexoses has been detected. Last, the cross-ring fragment at *m /z* 221 (C_6_ H_10_ O_5_ + C_2_ H_4_ O_2_) provides evidence that the Pent_4_ Hex chain and Pent_3_ Hex are branched on the inositol moiety *via* the hexose residue. Following the same reasoning, according to the fragmentation pattern obtained from candidates at *m /z* 716 and 724 (**Fig. 4A and 4B**), we attributed the respective fragments at m/z 895 and 911 to the presence of side chains composed of one hexose linked to the Hex-UA-Ino-P core and four pentoses and one additional deoxyhexose (m/z 716) or hexose (m/z 724). We finally deduced that the candidates at *m / z* 782 and 790 are characterized by one additional pentose (**Fig. 4C and 4D**).

**Figure 3.**
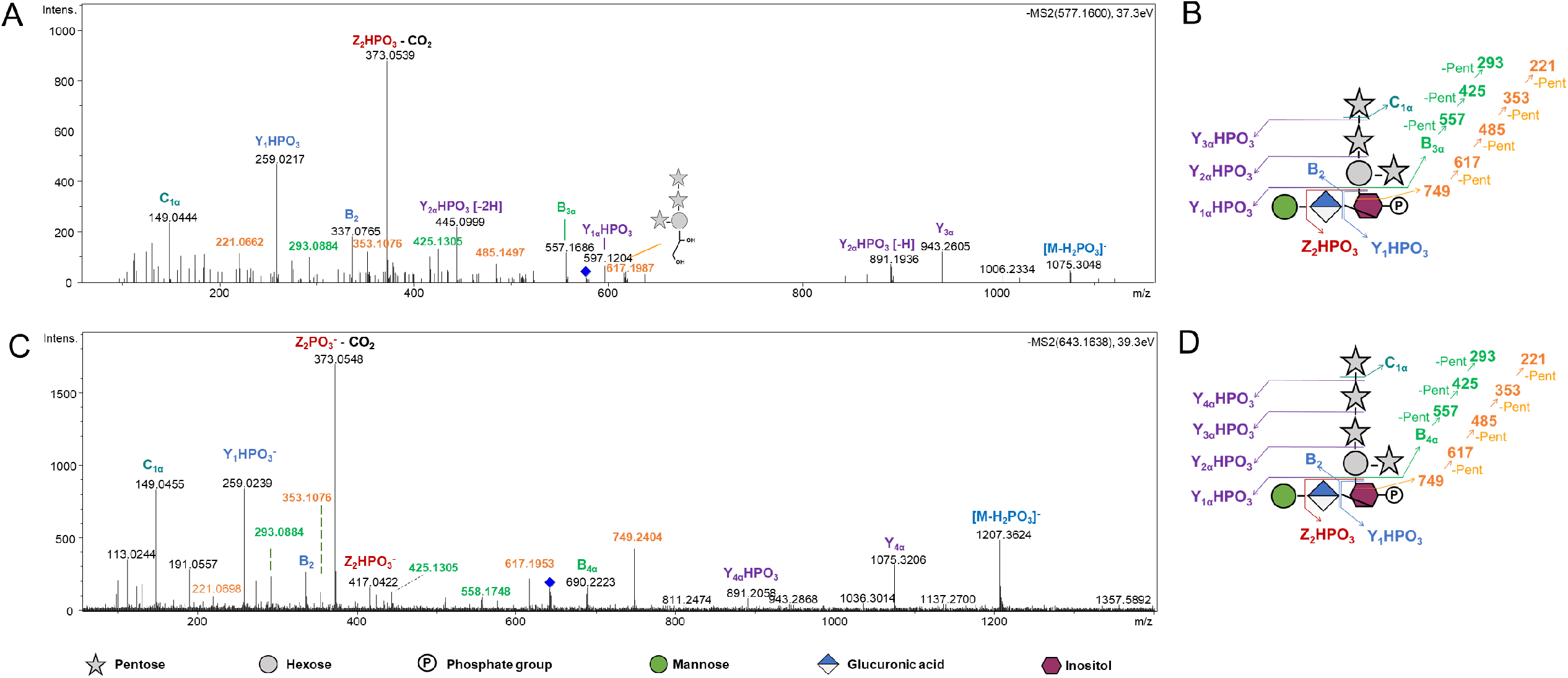
Substituted InositolPhosphate Glycans (I(P)Gs) are accumulated upon *Arabidopsis thaliana-Botrytis cinerea* infection. **A)** MS^2^ fragmentation pattern of *m/z* 577 in negative mode. **B)** Proposed fragmentation scheme for m/z 577. **C**) MS^2^ fragmentation pattern of *m/z* 643 in negative mode. **D**) Proposed fragmentation scheme for m/z 643. GlcA: Glucuronic acid, Intens.: signal intensity. Pent: Pentose

**Figure 4.**
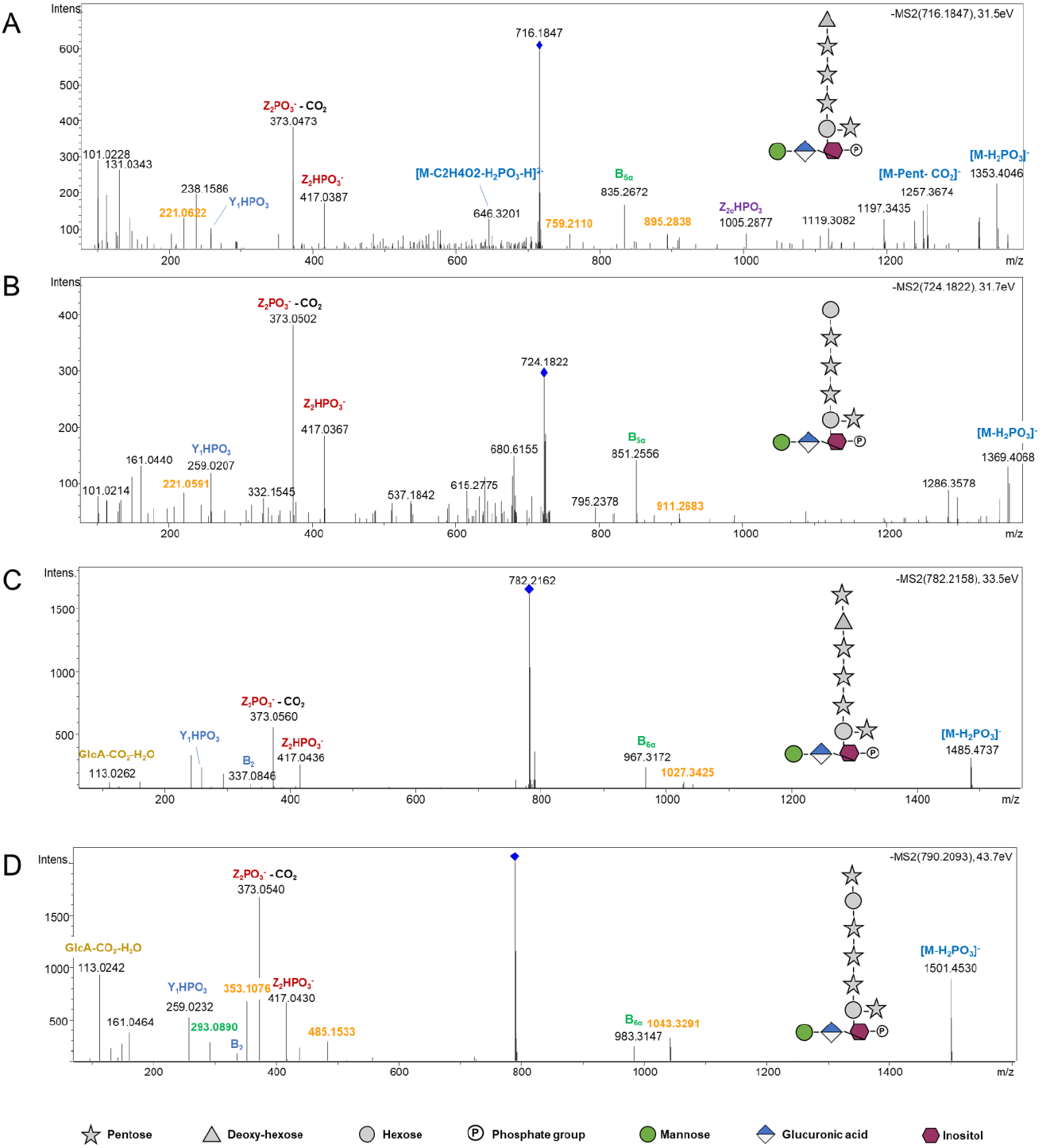
MS^2^ fragmentation pattern of *m/z*. **A**) 716, **B)** 724, **C)** 782 and **D)** 790 in negative mode. GlcA: Glucuronic acid, Intens.: signal intensity.

Therefore, we identified 7 inositolphosphate glycans (IPG) and one inositol glycan (IG) over-accumulated upon *A. thaliana / B. cinerea* interaction and present in cell wall water extract of healthy leaves.

In order to explore if these I(P)Gs could derive from glycosyl inositol phosphoryl ceramides (GIPCs), we analyzed the cell wall water extract of *gmt.1.3* (GIPC Mannosyl Transferase) mutant which is affected in GIPC glycan biosynthesis^14^. 14-day-old light-grown wild-type and *gmt1.3* plants that still displayed similar heights were analysed. In WT cell wall water extract, we detected six of the IPGs and IG accumulated upon infection (**Fig. 5A and B**). By contrast, none of the oligosaccharides described above were detected in the *gmt1* mutant. We found ions *at m / z* 435 (C_12_ H_20_ O_12_ P) and 355 (C_12_ H_20_ O_12_) absent in WT (**Fig. 5A and B**), that could correspond to GlcA-Ino-P and GlcA-Ino, respectively. The *m / z* 435 fragmentation gave rise to ions at 259 m/z, likely formed by the loss of an uronic acid, and at *m /z* 97 and 79 diagnostic of phosphorylated groups (**Fig. 5C**). The 355 fragmentation indicates the presence of a C_6_ H_10_ O_6_ moiety and an uronic acid absent in the WT sample that we attributed to GlcA-Ino (**Fig. 5D**). It is worth to note that we did not detect any ion corresponding to the GlcA-Ino core substituted by side chains questioning the GIPC glycan biosynthetic machinery. We concluded that the 7 I(P)Gs accumulated upon *A. thaliana /B. cinerea* interaction derive from Serie A and Series E to H GIPCs and would be released by GIPC phospholipase(s) C.

**Figure 5.**
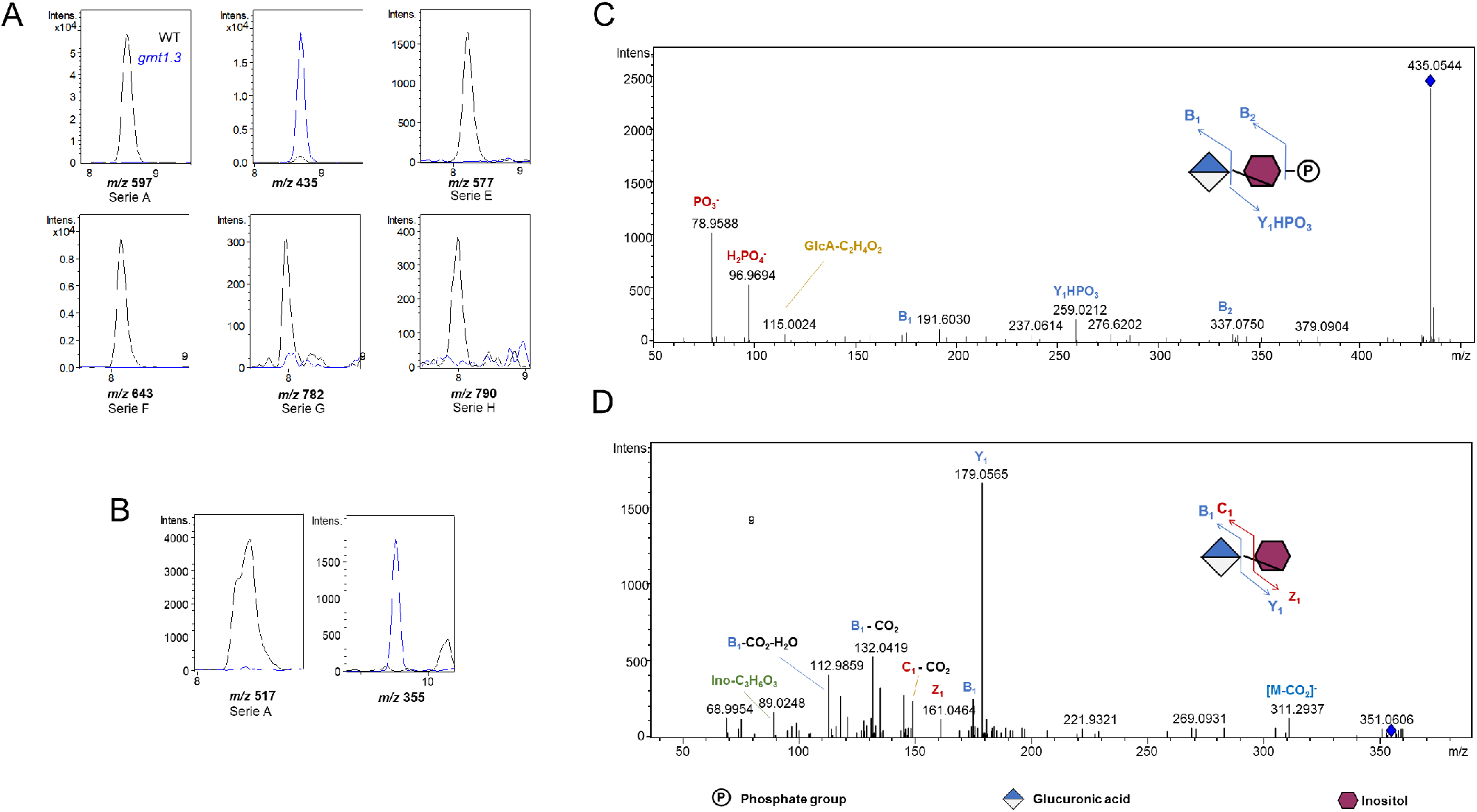
Inositol(Phosphate) Glycans (I(P)Gs) from cell wall water extract of *gmt1.3* mutant. Extracted ion chromatograms of IPG (**A**) and IG (**B**) in cell wall water extract of WT and *gmt1.3* 14*-*day-old plantlets. MS^2^ fragmentation pattern of m/z (**C**)435 and (**D**) 355. Intens.: signal intensity.

**Figure 6.**
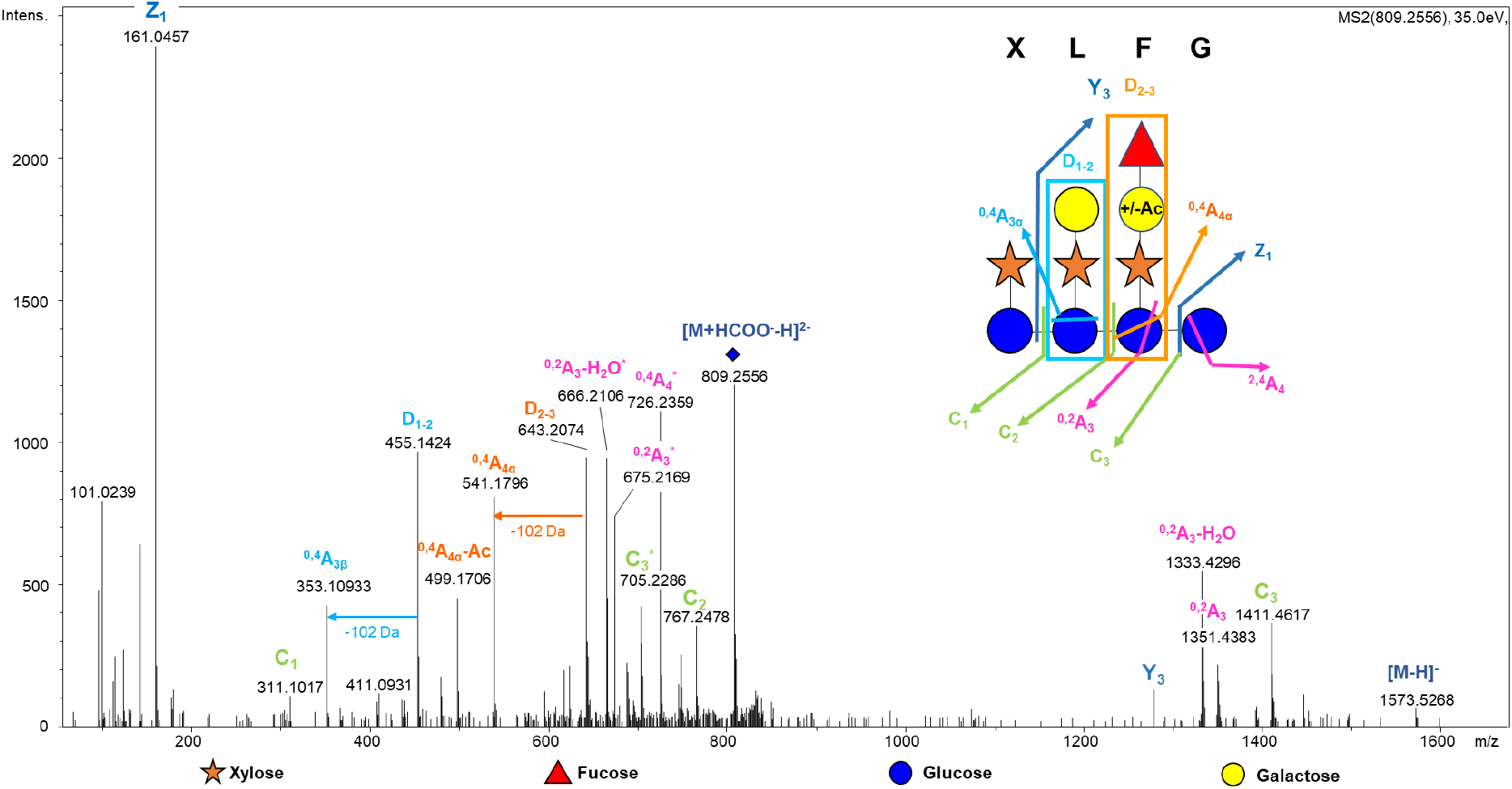
An acetylated and fucosylated xyloglucan oligosaccharide is accumulated upon *Arabidopsis thaliana-Botrytis cinerea* infection. MS^2^ fragmentation pattern of *m/z* 809 and a proposed fragmentation scheme.

The last candidate for which the software predicted the formula at 809 m/z is doubly charged (predicted formula C_59_ H_94_ O_51_ P^2-^) and detected as a formic acid adduct [M+HCOO^-^-H]^2-^. Its fragmentation gives rise to its [M-H]^-^ form at m/z 1573 and to ions at m/z 161 (anhydro-hexosyl unit), 1411 [M-C_6_ H_10_ O_5_ -H]^-^ and 1279 [M-C_6_ H_10_ O_5_ -C_5_ H_8_ O_4_ -H]^-^ suggesting the presence of hexosyl and pentosyl residues. We assigned the doubly charged ion at m/z 726, produced via a 120-Da loss to a ^2,4^A _n_cross-ring cleavage in a C_4_ -substituted hexose. Ions at 675 [M-Hex-C _2_H_4_ O_2_ - 2H]^2-^ and 666 [M-Hex-C_2_ H_4_ O_2_ -H_2_ O-2H]^2-^ were next assigned to a ^0,2^A_n-1_ cross-ring cleavage diagnostic of C_4_ - and C_6_ -substituted hexose. Last, the prominent and smaller single charged fragments at m/z 643 and 455 are accompanied by ions at m/z 541 and 353, respectively, which corresponds to a 102-Da neutral loss. This MS^2^ pattern is diagnostics of xyloglucan oligosaccharide fragmentations which are characterized by prominent D-type fragment ions. They result from double cleavage and correspond to entire inner xyloglucan side chains^15^. They produce in turns ^0,4^A_i_ cross-ring cleavage ions, diagnostic for the presence of (1,6)-linkage, via a 102-Da loss from anhydro-glycosyl unit. We therefore attributed the D-type ion at *m /z* 455 and its ^0,4^A_•_ corresponding ion at m/z 353 to the presence of Galactose-Xylose*-*Glucose block (represented by the letter L) and *m /z* 643 (D), 541 (^0,4^A_i•_) and 499 (^0,4^A_i•_ -Acetyl) to the presence of an acetylated Fucose-Galactose-Xylose*-* Glucose block (represented by the letter F). According to these results and to the well-known structure of xyloglucan, we concluded that the candidate at m/z 809 is an acetylated XLFG oligosaccharide.

In conclusion our study shows that structurally diverse oligosaccharides are produced *in planta*. Upon infection, *B. cinerea* secretes a xyloglucanase^16^, it is therefore consistent to detect xyloglucan fragment produced concomitantly with OGs. By contrast, the presence of I(P)G is far more surprising. The study of *gmt1.3* mutant affected in GIPC mannosylation showed that these I(P)Gs originate from GIPCs. The existence of an IG, originating from serie A GIPC, has already been described in extracellular space of culture cells during the period of rapid cell growth^17-18^. However, although in mammals IPGs were described over 30 years ago^19^, IPGs have not yet been identified in plants.

Here, we demonstrate that IPGs from serie A to H, present in cell wall extract of healthy plant are preferentially accumulated upon infection. This suggests that a GIPC phospholipase C activity which cleaves phospholipids before the phosphate group, is specifically promoted in that condition. It is worth to note that the non-specific phospholipase C4 (NPC4) has been shown to be involved in GIPC hydrolysis in response to phosphate deficiency^20^ and consequently, IPGs could act more broadly as stress-related molecules. The accumulation of serie A IG (m/z 517) also suggests that a GIPC phospholipase D, which cleaves after the phosphate moiety, could be activated. A phospholipase D activity has been detected in cabbage and in *A. thaliana* ^21^. This GIPC-PLD prefers GIPC containing two sugars^22^ which is consistent with the fact that Man-GlcA-Ino is the most highly produced in our conditions. However, we cannot rule out another scenario that would imply a phospholipase C activity and a subsequent phosphorylase. Finally, the I(P)G diversity observed questions the existence of diverse phospholipases C and D with different substrate preference. The identification of these enzymes, their impact in membrane properties as well as the role of such IPG and IG diversities need to be investigated.

The absence of series E to H IPGs lacking mannose residue in the *gmt1.3* mutant suggests that mannose adding is indispensable prior to the transfer of other sugars. More interestingly, this also questions the role of complex GIPCs in the *gmt1.3* phenotype. Indeed, this mutant *gmt1.3* displays a severe phenotype and a constitutive hypersensitive response^14^. The role of IPGs as signaling molecules needs therefore to be investigated in this mutant and in plants in general.

In mammals, IPGs are extracellularly-produced by insulin-sensitive cells in response to insulin treatment from glycosylphosphatidylinositol cleaved by phospholipase C^23^. They promote lipogenesis, glycogenesis^24^ and interestingly, are also important component of T-cell growth^25^. Insulin is involved in controlling blood glucose levels and its role in shaping immunity through modulating T cell metabolism is being more and more documented^26^. Taken together with the fact that I(P)Gs are accumulated upon infection in plants, we do believe that considering I(P)Gs as damage-associated molecular pattern and studying roles of I(P)Gs in defense will undoubtedly open new avenues of research.

## Material and methods

### Plant Material

Infection assays were performed on *A. thaliana* WT Wassilewskija (Ws-0) plants grown in soil in a growth chamber at 22 °C, 70% humidity, under irradiance of 100 •mol·m−2 ·s −1 with a photoperiod of 8-h light/16-h dark. Senescent *A. thaliana* WT (Ws-0) plants plants were grown in long-day conditions (16/8 h light/dark) in soil in a greenhouse. Seeds from *gmt1.3* mutants were a kind gift of J. Mortimer. *A. thaliana gmt1.3* and Colombia seeds were grown on 1/2 × MS media plates positioned vertically for 14 days under constant light at 23°C.

### Alcohol insoluble residue and cell wall water extract

Leaves and plantlets were grinded in five volume of ethanol 96 % and the supernatant was removed after centrifugation at 10,000*g* for 5 min. The pellet was washed with 70% ethanol with subsequent centrifugation until being uncolored and dried in a speed vacuum concentrator at room temperature. Two volumes of distilled water were added to the alcohol insoluble residue obtained and leave one hour at room temperature. The supernatant was collected after centrifugation at 10,000*g* for 5 min and dried in a speed vacuum concentrator at room temperature.

### Oligosaccharides accumulated upon *A. thaliana-B.cinerea* interaction

Oligosaccharides were produced and analyzed according to Voxeur et al., 2019. Briefly, the wild-type *B. cinerea* B05.10 strain was grown on potato dextrose agar at 23 °C under continuous light. After 10 days, the spores were washed from the surface of the plate using Gamborg’s B5 basal medium, 2% (w/v) fructose and 10 mM phosphate buffer. Fungal hyphae were removed by filtering. The concentration of spores was determined using a Malassez cell and adjusted to a final concentration of 3.10^5^ conidia/mL. Isolated *A. thaliana* leaves of 5-week-old plants were immersed in a *B. cinerea* suspension (6 leaves for 10 ml of suspension at 3 × 10^5^ spores/ml) and incubated on a rotary shaker at 100 rpm at 23 °C during 12, 15 and 18 h. The liquid medium was next collected and an equal volume of 96% ethanol was added. After centrifugation at 5000 g during 10 min, the supernatant was collected and dried in a speed vacuum concentrator at room temperature. The obtained pellet was then diluted. The equivalent of the digestate of 3 leaves of 5-week-old *A. thaliana* plants was dried and diluted in 200 μl. 10 μl were injected for MS analysis.

### Oligosaccharide analysis

Samples were diluted at 1 mg/ml in ammonium formate 50 mM, formic acid 0.1%. Chromatographic separation on high-performance size-exclusion chromatography was performed on an ACQUITY UPLC Protein BEH SEC Column (125Å, 1.7 µm, 4.6 mm X 300 mm, Waters Corporation, Milford, MA, USA). Elution was performed in 50 mM ammonium formate, formic acid 0.1% at a flow rate of 400 μl/min and a column oven temperature of 40 °C. The injection volume was set to 10 μl. MS-detection was performed in negative mode with the end plate offset set voltage to 500 V, capillary voltage to 4000 V, Nebulizer 40 psi, dry gas 8 l/min and dry temperature 180 °C.

Major peaks were annotated following accurate mass annotation, isotopic pattern and MS/MS analysis. The MS fragmentation pattern is indicated according to the nomenclature of Domon and Costello^27^. For the targeted analysis, the theoretical exact masses were used with 4 significant figures with a scan width of 5 ppm. The resulting extracted ion chromatograms were integrated.

### Data processing

The .d data files (Bruker Daltonics, Bremen, Germany) were converted to .mzXML format using the MSConvert software (ProteoWizard package 3.0^28^) mzXML data processing, mass detection, chromatogram building, deconvolution, sample alignment, and data export were performed using MZmine 2.52 software (http://mzmine.github.io/). Mass list were built using a retention time window of 6.0-8.0 min and next, we used the ADAP chromatogram builder^29^ with group size of scan 5, peak detection threshold of 800, a minimum highest intensity of 1500 and m/z tolerance of 0.01 m/z. We deconvoluted the data with the ADAP wavelets algorithm using the following setting: S/N threshold 8, peak duration range = 0.01-0.2 min RT wavelet range 0.02-0.1 min. Finally, we selected candidates with a surface area superior to 18000 at 18 hpi and results were deisotoped manually.

## Author Contributions

Performed the experiments: AV, JS. Analyzed the data: AV. Wrote the paper: AV, SV.

